# Paradoxical stabilization of relative position in moving frames

**DOI:** 10.1101/2021.01.23.427924

**Authors:** Mert Özkan, Stuart Anstis, Bernard M. ’t Hart, Mark Wexler, Patrick Cavanagh

**Affiliations:** Department of Psychological and Brain Sciences, Dartmouth College, Hanover, NH 03755, USA; Department of Psychology, University of California at San Diego, La Jolla, CA, USA; Centre for Vision Research, York University, Toronto, ON, Canada; INCC and CNRS, Université de Paris, Paris, France; Department of Psychology, Glendon College, Toronto, ON M4N 3M6, Canada

**Keywords:** vision, position, motion, visual constancy

## Abstract

To capture where things are and what they are doing, the visual system may extract the position and motion of each object relative to its surrounding frame of reference^e.g., 1,2^. Here we report a particularly powerful example where a paradoxical stabilization is produced by a moving frame. We first take a frame that moves left and right and we flash its right edge before, and its left edge after, the frame’s motion. For all frame displacements tested, the two edges are perceived as stabilized, with the left edge on the left and right edge on the right, separated by the frame’s width as if the frame were not moving. This illusory stabilization holds even when the frame travels farther than its width, reversing the actual spatial order of the two flashes. Despite this stabilization, the motion of the frame is still seen, albeit much reduced, and this hides the paradoxical standstill of relative positions. In a second experiment, two probes are flashed inside the frame at the *same* physical location before and after the frame moves. Despite being physically superimposed, the probes are perceived widely separated, again as if they were seen in the frame’s coordinates and the frame were stationary. This illusory separation is set by the distance of the frame’s travel, independently of its speed. This paradoxical stabilization suggests a link to visual constancy across eye movements where the displacement of the entire visual scene may act as a frame to stabilize the perception of relative locations.

## Introduction

Vision simplifies the dynamic world around us by coding motions and positions of objects relative to the frame that surrounds them^1^, up to and including the frame of the whole visual field. Studies have shown that frames and backgrounds have very powerful influences on vision, changing what we judge to be “up”^3,4^ and what direction we think is straight ahead^5,6^. When a frame is in motion, it can alter the impression of our own motion^7^ or that of an object within the frame^1,2,8,9^.

Here we report that a moving frame also triggers a paradoxical stabilization. When a frame is in motion for a second or less, and probes are flashed just before and after the motion, the separation between the probes is seen as if the frame were stationary. This stabilization is found for probes flashed on the edges of the frame (Experiment 1) or within the frame (Experiment 2) and it holds even though the edges of the frame are clearly seen to move (Movie 1, Fig. 1). These effects are the strongest illusions of position yet reported for steady gaze and we suggest that there are links between this paradoxical frame stabilization and visual constancy – our ability to see the world as stable despite the large shifts of the visual scene on our retinas every time we move our eyes.

**Figure 1.**
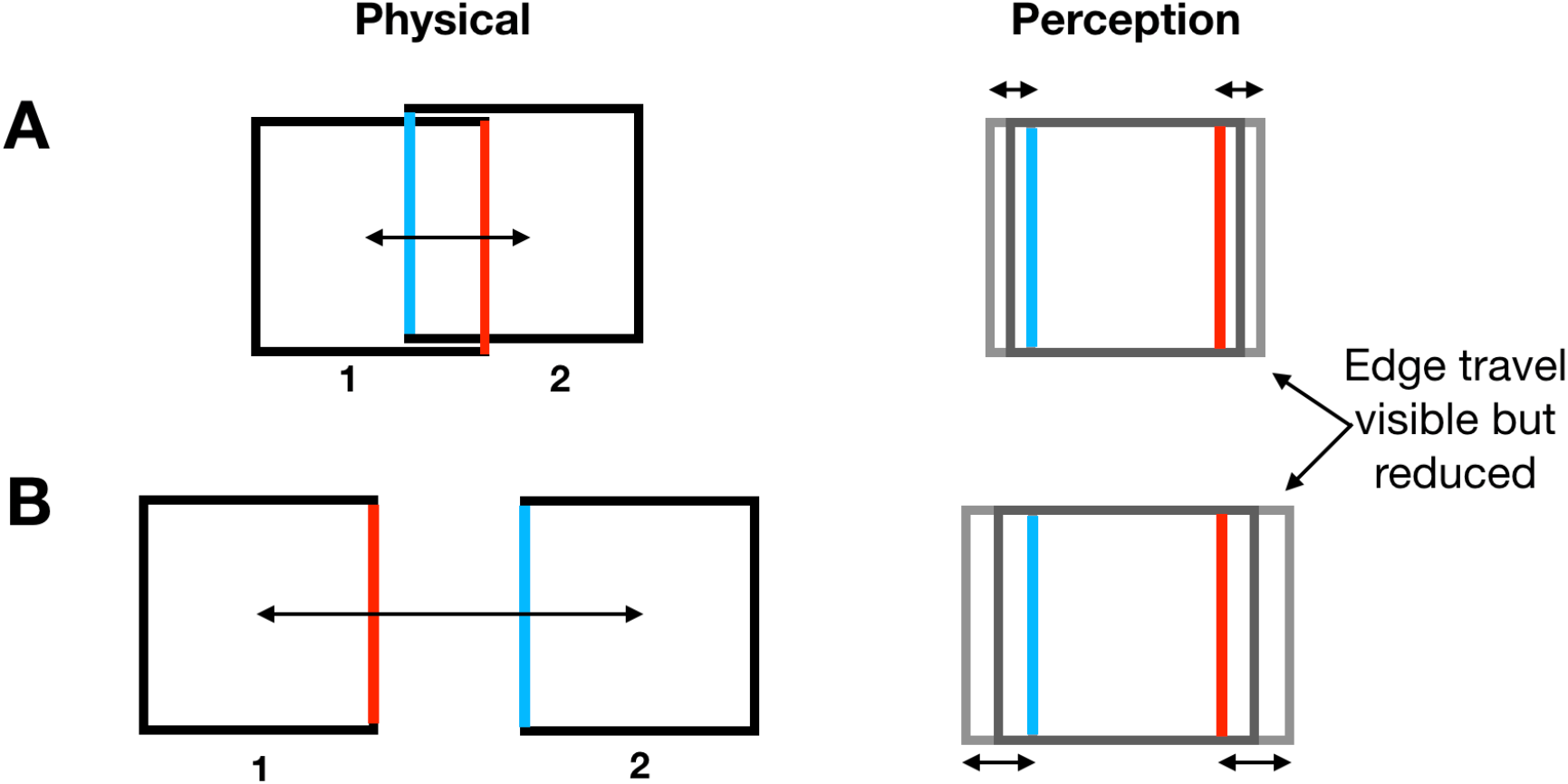
Paradoxical frame stabilization. **A)** The frame moves left and right by 2/3 of its width but instead of seeing the inter-edge spacing of 1/3 the frame’s width, as marked by the blue and red edges, the separation of the edges appears almost as large as the entire frame’s width – as if the frame were not moving. Paradoxically, the edges of the frame are still seen to move, although less so than their real travel. (The slight vertical offset of the frame is a graphical convenience, the frames had no vertical displacement in the experiments.) **B)** If the frame moves more than its width, the red edge is physically to the left of the blue and yet the blue still appears to the left of red, separated again by almost the width of the frame (Movie 1).

Our first experiment examines this paradoxical stabilization of relative position and the following experiments reveal equivalent effects of this stabilization on probes flashed within the frame.

## Results

### Experiment 1: Flashes at the edges of a moving frame are localized as if the frame were not moving

Participants report the perceived separation between parts of the frame flashed just before and just after the frame moves. In the first condition, the right edge of the frame flashed when it was the left end of its path, and then the left edge when the frame was at the right end of its path (Fig. 2, left panel). In the second condition, the left edge flashed at both the left and right ends of the frame’s path so that the physical separation between the flashes was equal to the frame’s travel (Fig. 2, right panel). Participants adjusted a pair of markers at the upper right of the display to indicate how far apart the flashed edges appeared. The results are clear and dramatic. As the physical separation between the flashed edges changed from 12.5° to −7.5°, the perceived separation remained relatively constant (Fig. 3A) at a value only slightly less than the 12.5° physical width of the square. Importantly, the same separation was reported even when the frame was virtually static (the baseline judgment—the left most data point). The measurement technique appears to underestimate the width of the frame so that the relatively constant setting at all path lengths indicates that the perceived separation was actually quite close to the perceived full width of the frame (average of 91.5% of baseline separation across the 4 non-baseline settings). Remarkably, this was true even when edges had reversed their relative positions (at the two longer path lengths). A one-way ANOVA showed no significant effect of path length on the perceived spacing (Greenhouse-Geisser corrected F(1.45, 11.31)=2.65, p=0.127). Despite this effective stabilization of relative positions, the movement of a single edge was clearly seen in condition 2 and did vary with the physical travel, being on average about 50% of the physical distance (Fig. 3B). The large but constant perceived inter-edge spacing in condition 1 is paradoxical: it would be expected only if the frame itself were perceived as stationary. But it is not – the perceived travel of the single edge showed that the motion of the frame was clearly visible, even if reduced. It is as if the locations of the flashed edges are reported in the coordinate system of the frame as if it were stationary although the frame’s movement is quite apparent (but underestimated).

**Figure 2:**
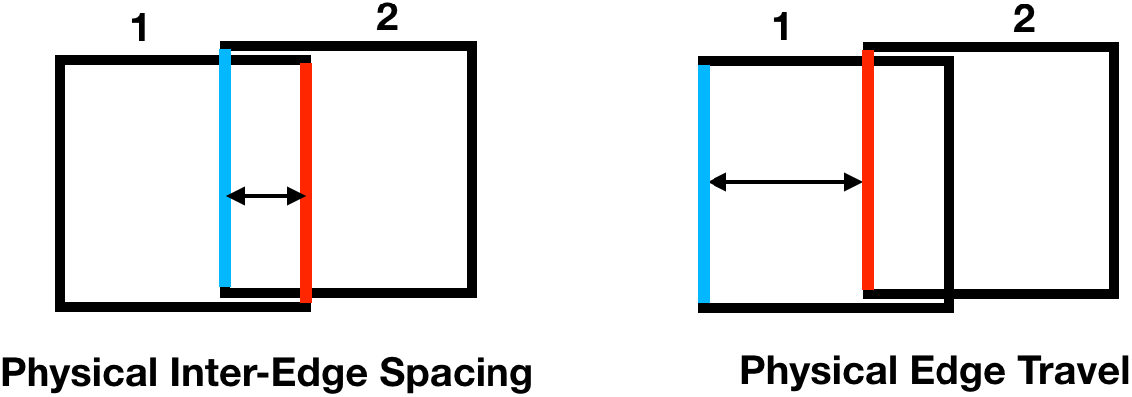
Measuring perceived frame travel. **Left.** In the first condition, opposite edges of the frame flash at the two ends of the travel as in Fig. 1A. Participants adjusted a pair of markers at the upper right of the display to indicate how far apart the flashed edges appeared. **Right.** In the second condition, the left edge is flashed at the two ends of travel and the space between them is the distance the frame travels. Participants again adjusted markers to indicate the perceived spacing.

**Figure 3:**
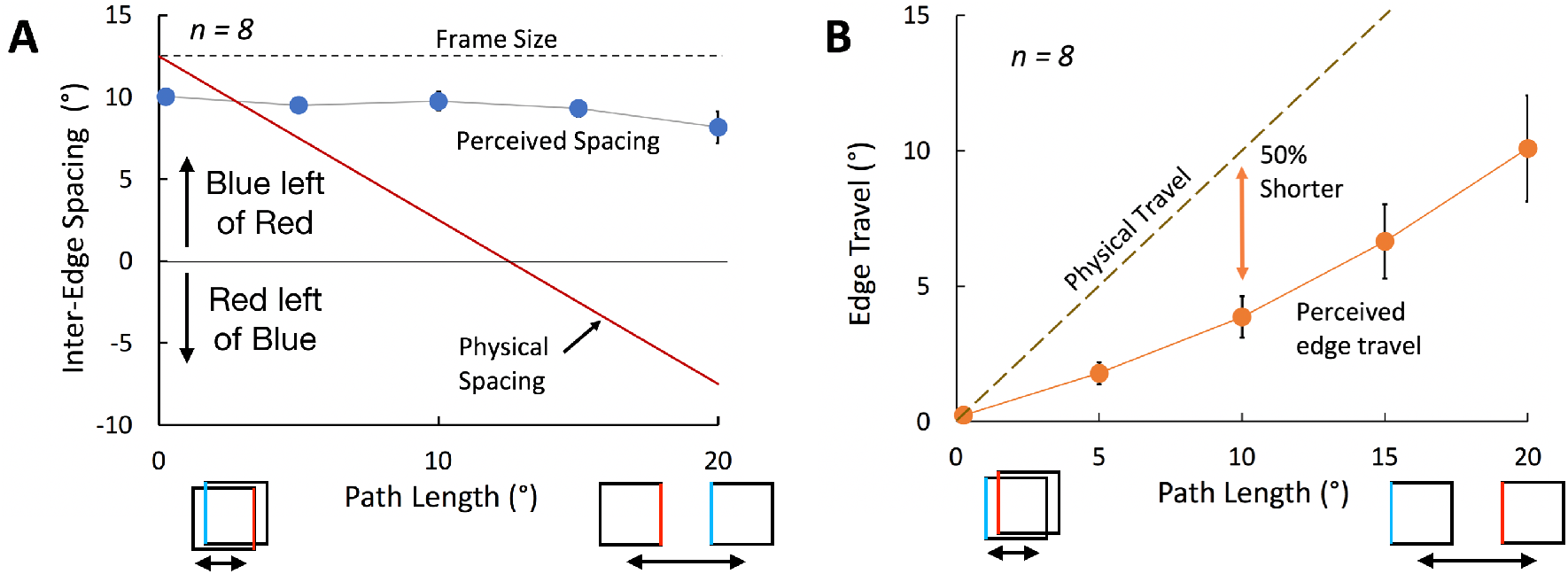
**A) Inter-edge spacing.** The perceived distance between the blue and red flashed edges remains positive (blue left of red) and constant even when it is physically negative (where red is actually left of blue). The left most data point is the baseline judgment of the frame’s width when the frame is virtually stationary. **B) Single edge travel.** The frame’s perceived motion (orange symbols) is about 50% of its physical motion. Error bars show ±1 S.E. where larger than the data symbols.

### Experiment 2: Stabilization-induced position shifts

Here we examine whether probes flashed inside the frame are also perceived in the coordinate system of the paradoxically stabilized frame, shifted far from their physical locations. In this experiment (Fig. 4, Movie 2) a frame moved left and right while two targets were flashed at the *same* physical location on the screen every time the motion reversed direction. The first flashed when the frame was at the left end of its travel, so that the flash was close to the right side of the frame. The second flashed when the frame was at the right end of its travel placing the same flash location close to the left side of the frame (Fig. 4). As before, participants adjusted a pair of markers at the upper right of the display to indicate how far apart the flashes appeared. The first condition tested the perceived separation of the flashes over a 64-fold variation of the frame’s speed with the frame size set at 15° and the path length at 10°. Despite being physically superimposed, the two flashes appeared to be shifted away from each other by about the distance the frame had traveled (Fig. 5A). There was no significant effect of speed on the perceived shift (Greenhouse-Geisser corrected F(1.47, 18.71)=2.013, p=0.19), and the average illusory shift was 96% of the path length across the 6 speeds. The flashes were therefore seen in their location in the frame as if the frame were almost stationary! Surprisingly, this stabilization held up even at the slowest speed we tested (a bit more than 1 second for each traverse) but clearly the effect must drop off quickly at even slower speeds like that seen at the beginning of Movie 1 where each traverse lasted 1.5 seconds and there appears to be no stabilization.

**Figure 4:**
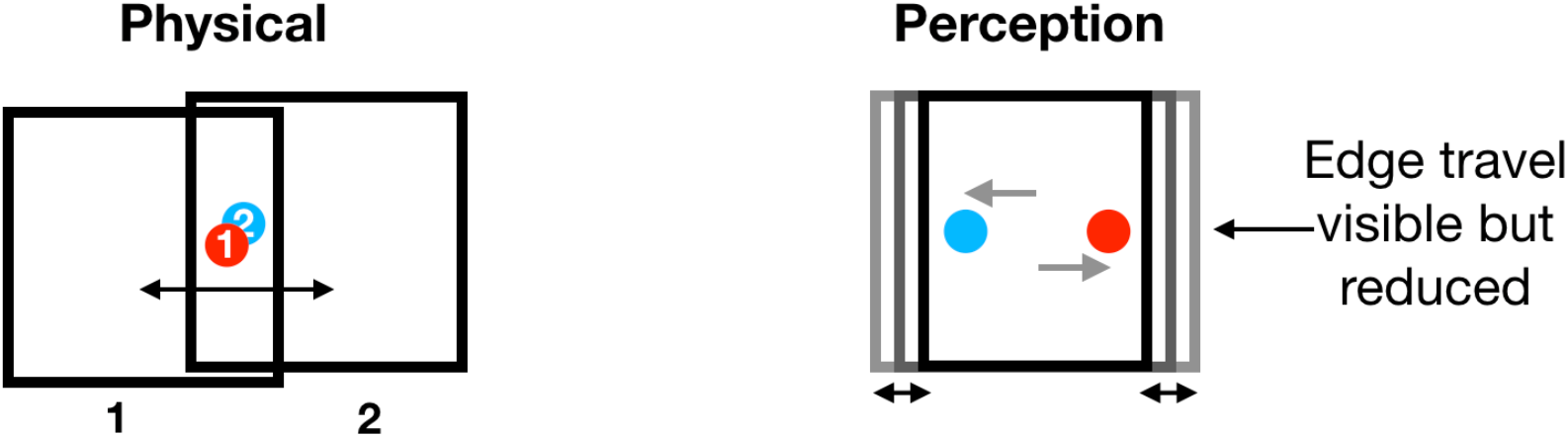
Probes flashed within the frame. Flashed probes are seen at their locations relative to the frame, as if it was almost stationary, shifted far from their physical locations (Movie 2).

**Figure 5:**
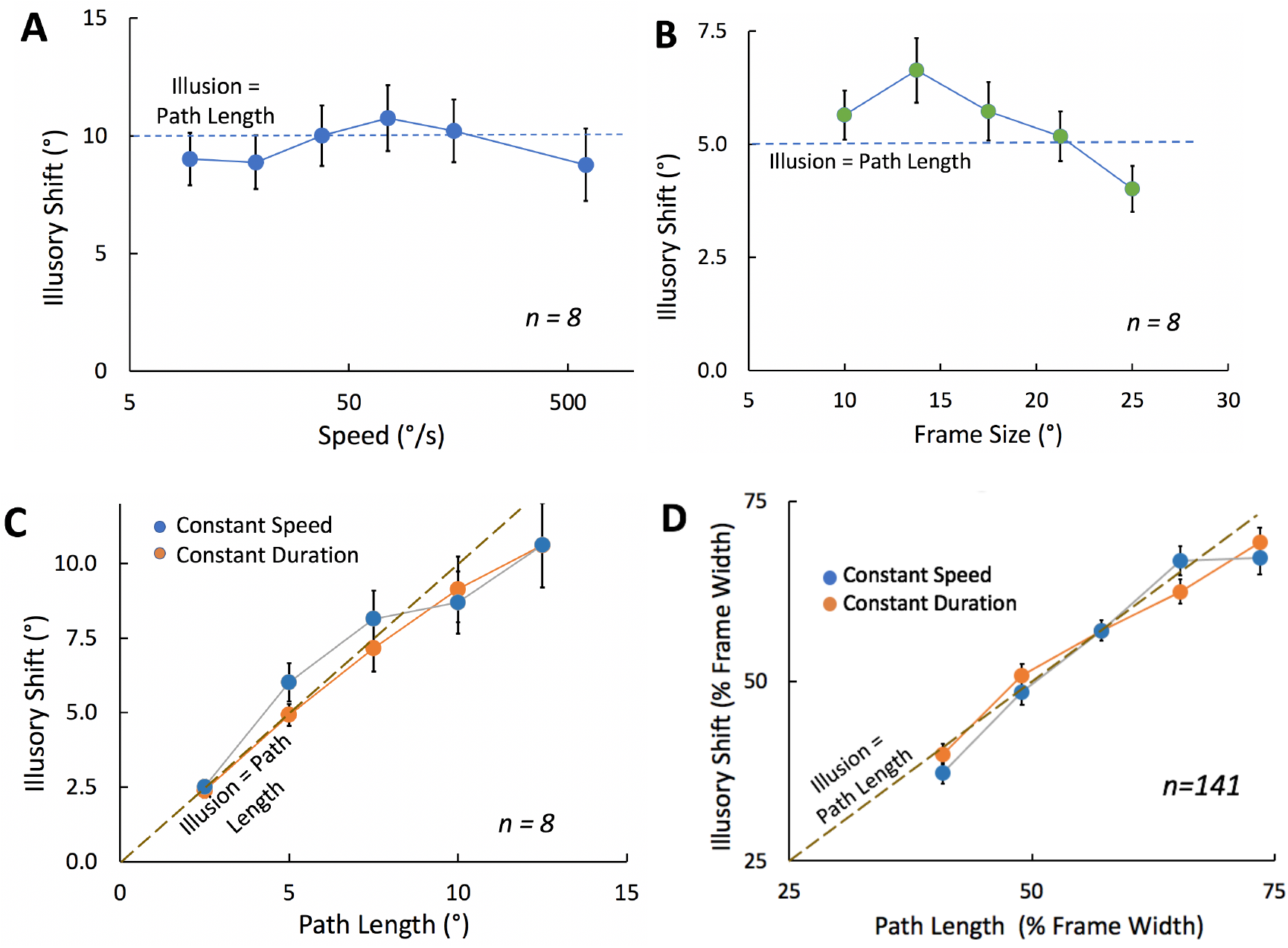
**A)** Illusory shift as a function of speed (on a log scale). Path length was 10°. **B)** Illusory shift as a function of frame size. Path length was 5°. **C)** Illusory shift as a function of path length for constant speed or constant duration motion. **D)** Same as C but for 141 participants run on-line. In all cases, error bars show ±1 S.E. when larger than the data symbols.

In the second condition, the speed and the frame’s path length were fixed at 30°/s and 5°, respectively, while the size of the frame varied from 10° to 25°. There was significant variation with size with the perceived shift dropping at larger frame sizes (Fig. 5B, F(4,28)=11.17, p=0.000015) perhaps because the frame was getting close to the borders of the monitor. Nevertheless, overall, the apparent offset between the flashes was close to the path length – the average across the 5 frame sizes was 109% of the path length.

In the third condition, the frame size was fixed at 15° while the path varied from 2.5° to 12.5°. On half the trials, the speed was held constant at 30°/s while on the other half, the duration of movement was held constant at 500 ms. The settings (Fig. 5C) show that the perceived offset of the flashes increased relatively linearly with the path length of the frame and did so similarly whether the speed was constant, or the duration of travel was constant. The mean setting was slightly but significantly higher for constant speed than constant duration (F(1,7) = 14.39, p=0.007) and there was a significant interaction (F(4,28) = 6.81, p=0.001). However, these two effects accounted for only 0.4% and 1.0% of the total variance (eta-squared) whereas the variation in path length accounted for 92.1%. Averaged across the 5 path lengths, the illusory shift was 97% of path length.

We were able to repeat this last condition as an online experiment with York University undergraduates (see demonstration here https://run.pavlovia.org/mthart/pubdemo/html/), as part of a battery of cognitive tests. The monitor size and viewing distance varied from participant to participant and these values were available for about 50% of them. The results for these participants showed that the size of the frame in degrees of visual angle did not affect the outcome significantly so we averaged the results across all participants. The frame’s travel and the perceived separation are reported as percent of the frame size (Fig. 5D). The effects for the 141 participants who remained after screening (see methods in Supplemental Materials) were quite similar to those for the 8 in-person participants. Perceived offset was again determined by the frame’s path length and did not differ between the constant speed and constant duration conditions. The illusory shift was 97% of the path length (averaged across path lengths).

## Conclusions

Surprisingly, our data show that when a frame moves by less than about twice its width at a moderate to rapid speed, the relative distances between probes flashed before and after the motion are seen as if the frame were almost stationary. How can this be? The frame appears to be moving quite well — nobody reports that the frames are motionless. It is only our data that reveal that the frame’s displacement is discounted for the judgments of the separation between the flashes. At the same time, the measurements of a single edge of the frame do show that the frame is seen to move, even though less than its physical travel. This produces the impression of robust motion, hiding the paradoxical stability of relative positions.

### Why are relative positions stabilized?

The effect we report here is not simply a reorganization of the visual elements in frame-specific coordinates as proposed by Johansson in 1950^1^ and others^2,8^. The critical difference is that, in Johansson’s displays, the elements within a moving frame are seen to move relative to the frame, but they also are seen to share the common motion of the frame. Imagine someone waving to you from the window of a passing train, we see their hand waving up and down – its motion relative to the window – rather the sine wave it actually traces out. But we also see the person and their hand moving with the train, the motion they all have in common. In contrast, in our examples, the flashes escape the common motion – the displacement of the frame has been discounted and they have been effectively stabilized in world coordinates. We have no mechanistic explanation for this effect although it could be a consequence of the extraction of the frame-relative positions as suggested by Johansson^1^. Perhaps the flashed probes do not seem to move along with the frame because they occur only before and after the motion. In contrast, probes in Johansson’s displays were always presented during the frame’s motion. Our frame stabilization may also be a small-scale application of a more general processes that stabilizes the visual field when the eyes move – more on this below.

### What can act as a frame?

Here we use an outline square, but we have also found similar results with a moving cloud of random dots where even a few dots are enough to trigger the position shift of the probes^10^. This suggests that anything that moves as a group probably engages motion discounting and the paradoxical stabilization it produces. Further studies could clarify what qualifies as a group and also what transformations (e.g., translation, expansion / contraction, warping) produce stabilization.

### How does this relate to other frame effects?

There have been many demonstrations of the effects of frames on position, orientation, and motion decomposition, but none with effects of the magnitude we report here. The illusory displacements of almost 100% of the frame’s travel here is far stronger than the original motion induction effects of Duncker^2^ and others (review^11^) where the stationary probe was presented continuously during the motion. Possibly Duncker’s induced movement is reduced by the continuing evidence that the target is stationary during the frame’s motion. In contrast, our flashes were brief and, importantly, they were not present during the motion. Moreover, Duncker’s effect becomes noticeable only when the frame’s motion is very slow, between motion threshold and maybe three times motion threshold^12^. These speeds are much too slow to have any relevance to the abrupt displacement of the visual field that occurs with saccades. The frame effect that we see here is reliably large even for very abrupt displacements – in Fig. 3A, the right most data point is for a single jump of the frame with no intermediate positions.

### Eye movement confounds

One alternative explanation of the frame stabilization is tracking eye movements. If the eyes were to smoothly pursue the frame, it would be more or less stable on the retina and flashes at the same location on the screen will fall on very different locations on the retina. This might lead to a stabilization of the frame and a perceived shift of the flashes. However, the everyday effects of pursuit are quite different – a tracked object does not appear to be stabilized, and the positions of flashes during pursuit are also compensated for the pursuit motion — although not completely^13^. Whatever the case, it is easy to verify that the effects reported here persist with fixation (see Movie 1 or 2 and fixate a corner of the movie or another landmark on or near the display) and that they are no different with free viewing or with actual pursuit of the frame. Steady fixation on the flashed probe may reduce the effect for some viewers, and an actual fixation point near the discs will eliminate the frame’s effect, probably by giving high certainty information about the flash location.

### Relation to motion-induced position shifts

How are these frame effects related to motion-induced position shifts such as the flash lag^14,15^, flash grab^16^ or the shortening of motion paths^17^? In all these cases, motion also causes a shift in perceived position. However, there are several differences. First, the frame effects reported here are fairly constant across speeds up to quite high values (Experiments 2), whereas motion-induced position shifts for the flash lag^18^ and for the flash grab^16^ are speed dependent. Second, the perceived length of an oscillating motion path is shortened to about 70% of its true length^17^, perhaps due to position averaging^19^. We also find the frame’s path shortened here (Experiment 1) – but to 50% of its actual value, almost twice the effect. More importantly, even this shortening could not explain the perceived stability of relative positions which was equivalent to a 100% path shortening – a full discounting of the frame’s motion. Third, the frame stabilization had relatively global effects on position (Fig. 5B) – it was not greatly affected by the distance between the contours and the flash – in contrast, motion-induced position shifts like the flash grab decrease rapidly as the test moves away from the moving contour^16^. So, despite the resemblance of the motion and frame induced effects, there are qualitative differences that distinguish the two. It is likely that some stimuli^20^ may trigger both frame stabilization and motion effects and further studies will be required to clearly understand whether the two are related and how they are different.

### Visual constancy

Are the effects of a moving frame linked in any way to visual constancy, where the eyes move but the scene appears stable? Our results suggest a plausible, partial link. There have been many proposals to explain visual constancy, some that rely on extraretinal information, such as the assumption of a stable world^21,22^, the subtraction of efference copy^23,24^, or the remapping or updating of attended or landmark targets^25,26,27,28, 29^. Others suggest that purely retinal information is sufficient based on the discounting of common motion or displacement^30^. The stabilization of moving frames reported here falls in this second group of mechanisms, as the subtraction of the frame’s displacement is based solely on retinal signals. However, this falls short of an explanation of visual constancy for two obvious reasons. The first is that we found that the motion of the frame itself was not discounted – it was clearly seen to move even if its amplitude was reduced (Fig. 3B). The second is that retinal motion on its own was never an adequate explanation for visual constancy. Specifically, stabilization is not seen for retinal motion in the absence of eye movement commands. When pushing your eyeball or viewing a scene in a moving mirror or an unsteady video (review^31^), the visual scene clearly moves. Moreover, if an eye movement is attempted but produces no retinal motion due to afterimages^32^, paralyzing the extra-ocular muscles^32,33^, or optical stabilization^34^, the world nevertheless seems to move. Clearly, extraretinal signals are necessary for visual constancy; nevertheless, the robust frame stabilization reported here suggest that the frame effect would be an effective contributor, along with extraretinal signals, to constancy processes.

### Simulated saccades

The discounting of frame motion has been reported previously in simulated saccade studies^35,36,37,38,39,40^,. In these experiments, the display area, usually a reference ruler, is quickly shifted on a monitor or with a mirror. Probes presented at least 50 to 100 ms before the displacement are reported to be at their original location in the frame (e.g. retaining their ruler coordinates). However, because of the measurement technique, there is some doubt about whether the frame motion is discounted: the participants may have reported where they remembered the flash on the ruler at the time it flashed, well before the ruler moved. In contrast, our report technique avoids the ambiguity of the moving reference ruler and our results show that the separation between probes flashed before and after the motion is stabilized in world coordinates.

To conclude, we have demonstrated that when frames move, their displacements may be strongly suppressed, but more strikingly, probes in or on the frame that are flashed just before and after the frame’s movement are seen in the frame coordinates as if the frame were stationary: the frame’s motion between the first and second flash is largely discounted in perceiving the separation between the flashes. We suggest that this identifies a mechanism that contributes in part to the compensation for eye movements where the frame is the entire visual field.

## Acknowledgments

NSERC Canada (PC), VISTA (PC), Dartmouth PBS (PC), ANR (MW), UCSD Pathways to Retirement (SA). We would like to thank Don Macleod for helpful comments.

## Methods

### In-person Experiments

#### Participants

Eight individuals, including two of the authors, participated in the in-person experiments of this study (1 female; age range: 22-75, mean age = 46±7.4). All participants other than the two authors were naive to the purpose of this study and had normal or corrected-to-normal vision. Written, informed consent, approved by the York University HPRC and the Committee for the Protection of Human Subjects at Dartmouth College, was obtained from each participant prior to their experimental sessions.

#### Apparatus

All stimuli were generated an Apple Macintosh G4 computer with custom software written in C using the Vision Shell Graphics Libraries (Comtois, 2003). The display was presented on an LCD monitor with 60 Hz refresh rate and resolution of 800 x 600 pixels. The size of the display area was 40° x 30° (degrees of visual angle). Response adjustments were made with a track pad or mouse. Head movements were restrained with a chin rest and the viewing distance was 57 cm.

#### Stimuli

The screen was filled with a uniform mid-grey background while the lighter square frame had 50% contrast (Michelson) with the background. The square’s size varied depending on the experiment, but its contour always subtended 0.6°. The frame’s motion path was centered horizontally on the display and the vertical center of the frame was 3.75° below the display’s vertical midpoint to provide space for the measurement markers at the top right. Probes were alternately red and blue. In the first experiment, the probes were superimposed on the left or right contour of the frame and had the same width and height as the contour. In the second and third experiments, the two flashed probes were discs of 1.5° in diameter. In the continuous motion condition of experiment 3, the single probe was red and was also 1.5 ° in diameter. In all conditions, two adjustment markers were present in the upper right of the display, 10° horizontally from the screen center and 16.5° above the midpoint. In Experiment 1, the markers were red and blue vertical bars 0.6° wide and 1.5° tall; in Experiment 2, they were discs with the same size and color as the probes.

#### Procedure

Common to both experiments, each trial began with a beep following which the frame was present continuously and moved repeatedly back and forth horizontally. With one exception, the red and blue probes flashed alternately each time the motion reversed direction. They were presented for 33 ms, centered in the pause of the frame’s motion at the end of each transit. The pause at the end of the motion was adjusted to avoid the deterioration of the frame’s effect seen for short intervals between the repeating flashes (Cavanagh, Wexler, & Anstis, 2019). In any condition where the motion duration was less than 333 ms, the pause was increased from 33 ms (default value) to keep the inter-flash interval at 366 ms. Participants were instructed to look around the display wherever they wanted as they made their setting but to avoid fixating directly on a probe. Using a mouse or a trackpad, they adjusted the two markers at the top right until their separation matched that of the flashed discs (or the angle of the continuous path). When they were satisfied with their match, they pressed the space bar, and the next trial began. There were 4 repetitions of each trial. The responses were self-paced, participants could take a break at any time. The two experiments were run separately, and the sessions lasted about 10 and 20 minutes, respectively.

##### Experiment 1 Frame edges. Left vs right edge condition

The frame was 12.5° across, the motion duration was 166 ms and the pause was 200 ms. The path length took 1 of 5 values: 0.25°, 5°, 10°, 15°, and 20°. The shortest value was given a small offset so that it had some reversal transient. When the frame was at the left end of the trajectory, its right side briefly flashed red; when the frame reached the right end of the trajectory, its left side briefly flashed blue. Participants adjusted the two markers to indicate the separation they saw between the red and blue edges. Note that there was no overlap between frame at its leftmost and rightmost positions for the path lengths of 15° and 20° so the red right edge and blue left edges of the frame reversed locations in screen coordinates. *Left edge condition*. Same as the first condition except that it was the left edge that flashed at both ends of the frame’s travel. It flashed red at the leftmost end and red at the rightmost end. All the trials for the 2 conditions were randomly intermixed for a total of (10 × 4) 40 trials.

##### Experiment 2 Flashed discs. Speed condition

The frame was 15° across and the path length was 10°. The motion took 1 of 6 durations: 16 ms, 66 ms, 133 ms, 266 ms, 533 ms, and 1066 ms. The motion paused at each end of the travel for at least 33 ms and the flash of 33 ms was centered in that pause. The pause durations for the 6 motion durations were 350 ms, 300 ms, 233 ms, 100 ms, 33 ms, and 33 ms, respectively. *Size condition*. The path length was 5° and the motion duration was 166 ms with a pause of 200 ms at each reversal. The frame size took one of 5 values: 10°, 13.75°, 17.5°, 21.25°, and 25°. *Path length condition*. The frame was 15° in size. Path length took 1 of 5 values: 2.5°, 5°, 7.5°, 10°, and 12.5°. On half the trials, the motion duration was fixed at 500 ms with a 33 ms pause so that the speed increased as the path length increased. On the other half of the trials, the motion duration was adjusted to keep the speed of the moving frame constant at 30°/s, so 83 ms, 166 ms, 250 ms, 333 ms, 416 ms, with pauses of 283 ms, 200 ms, 33 ms, 33 ms, and 33 ms, respectively. All the trials for the 3 conditions were randomly intermixed for a total of (21 × 4) 84 trials.

**Figure S1.**
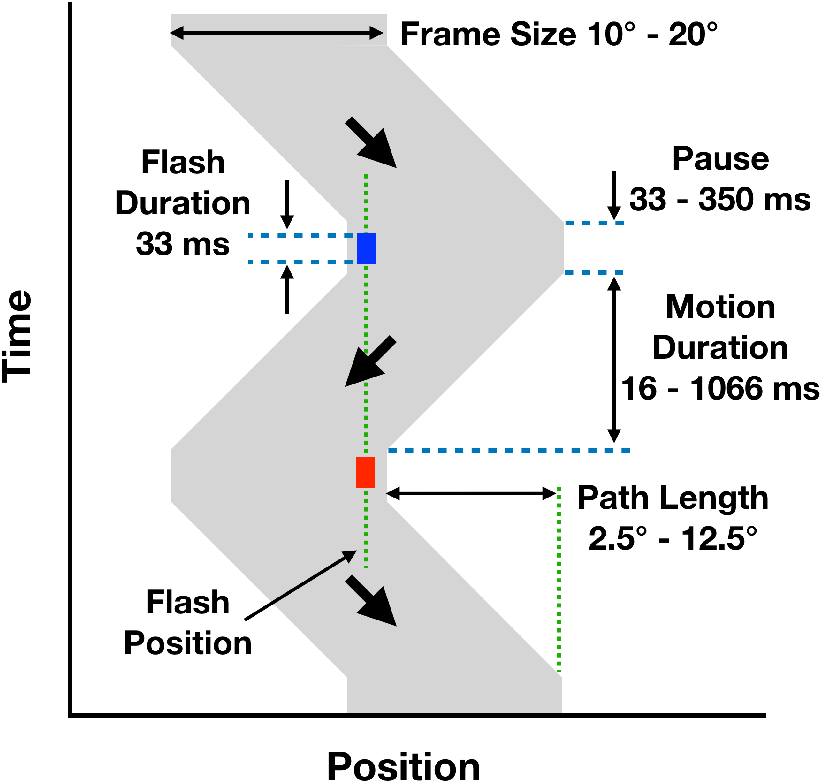
Timing for Experiment 2. The frame moved left and right. The motion paused at each reversal of direction, during the pause, the flashed probe was presented for 33 ms. The two flashes always had the same physical location. The pause had a minimum duration that was the same as the flash, 33 ms, but was longer whenever necessary to maintain a minimum of 366 ms between the onset of the two flashes.

#### Analysis

The 4 settings for each condition were averaged and then the mean and standard error of the mean across participants were calculated and plotted in Figs. 3 and 5. When standard error for a participant was high, a second session was run. The participant means for each condition were entered in the one-way and two-way ANOVAs reported in the text and available at https://osf.io/y5xrn/wiki/home/.

### Online Experiment

#### Participants

Participants were recruited from the York University population, and provided prior, written, informed consent. We selected only participants with self-reported normal or corrected-to-normal vision. All procedures were approved by the York University Human Participant Review Committee. We applied several inclusion criteria (see below). There were 274 participants with a complete data set and 141 remained following screening. The mean age was 22.1 ± 6.2 years (range: 17-50) including 106 females; 121 were right-handed, 15 left-handed, and 5 other.

#### Apparatus

The task was created in PsychoPy Builder (Pierce et al., 2019) and ran on the Pavlovia platform (https://pavlovia.org/). It was accessed from a Qualtrics questionnaire, embedded with other tasks as well as questions about the participant, the device used, and the presence of any surrounding distractions. We selected only those who indicated using a laptop or a PC (i.e. no phone or tablet), and who reported using a mouse or trackpad for the task (no touch screen), and who did not report any major disruptions during the task. Since each person used a different device, the screen size and viewing distance was variable. We asked people to measure the distance from the bridge of their nose to the center of the screen and to adjust an on-screen rectangle to a credit card pressed to the screen. Not all of the participants were able to do this, but for those who did (N=59), we calculated the size of the stimuli in degrees visual angle. An analysis of the perceived shift as a function of viewing angle across these participants showed that the effects were not influenced by the visual angle of the stimulus allowing us to average results across all participants.

#### Stimuli

On the bottom of the participant’s browser window were instructions that remained throughout the session and the number of completed trials over the total number of trials was shown at the top left of the screen. On the left were a pair of discs, one red, one blue, for adjustment that were also presented continuously. On the right side, the frame moved left and right and red and blue probes were flashed at each reversal of direction. See demonstration version here https://run.pavlovia.org/mthart/pubdemo/html/. The frame was a square whose height and width were 24.5% of the smaller of the height or width of the browser window, running in full screen mode. All further spatial values are given in percent of the frame width. The width of the frame’s contour was 8% of the frame size. The diameter of the adjustment and the flashed discs was 10.2% of the frame’s width. The separation of the two discs on the left could be controlled by moving their mouse or track pad left and right to indicate the perceived distance between the two flashed discs on the right. All 214 participants’ operating systems reported a refresh rate of 60 Hz and the dots were flashed for a duration that should ensure 2 frames (67 ms) with a 30 Hz refresh rate, so 4 frames (67 ms) for the 60 Hz refresh. Four catch trials were included with probes alone and no frame.

#### Procedure

There were exactly 22 trials in the experiment. In 12 of the trials, the probes were flashed at the same location. There were again 5 different path lengths: 41%, 49%, 57%, 65%, and 73% of the frame’s width combined with the two conditions: constant speed and constant duration. In the constant speed case, the motion duration increased in step with the path length (500 ms, 600 ms, 700 ms, 800 ms, and 900 ms, respectively) making the frame’s speed a constant 82% of the frame’s width per second. In the constant duration case, the duration was 700 ms throughout, resulting in 5 different speeds. There was one trial for each combination, except for 57% and 700 ms, present in both conditions, which had two trials in each set. There were an additional 6 stimuli with a vertical offset of 61% of the frame’s width between the two flashed probes. Three path lengths for the frame’s movement—41%, 57% and 73% of the frame’s width—were combined in both constant speed and constant duration trials, one of each. For constant speed, the motion duration increasing with the horizontal offset to give a constant horizontal velocity of 82% of the frame’s width per second. For the 3 constant duration trials, there was a fixed 700 ms motion duration. These data are not reported but are available with the rest of the data on the OSF site (see below). Finally, there were 4 catch trials where the probes were flashed with no frame present. Two of these had no vertical offset but a horizontal offset of 61% of the absent frame’s width, one with the red probe on the left, one with the red probe on the right. Two had only a vertical offset of 61% of the absent frame’s width and no horizontal offset one with red on top and one with blue on top. These data were intended to screen participants for accuracy but all responses were reasonable.

The 22 stimuli were ordered randomly for each participant, with a 1 s inter-trial interval. Participant were asked to change the horizontal position of the two continuously present dots on the left of the screen so that their horizontal offset (as well as angle in some stimuli) matched that of the two flashed dots on the right. When they were satisfied with their setting, they pressed the space bar, ending the trial and starting the next. It took participants on average about 6.5 minutes to complete the task.

Given the nature of online experiments, participants may not have been fully focused on the task, or may have misunderstood the short, written instructions. Based on the self-report criteria described above, 33 participants were removed for reports such as “just clicked as quickly as possible,” “didn’t understand the instructions,” “interrupted during test.” Of the remaining 241, 100 participants were removed for not meeting performance criteria. To eliminate outliers who may have been pressing the space bar without making adjustments we required that settings on the 12 main trials had to all be less than 3 times the maximum frame movement, and that a maximum of two settings could be 0. Of the original 274 participants, 141 remained.

#### Analysis

We used descriptive statistics (means and standard errors) to plot the effects of the frame’s five different movement amplitudes and the two main conditions (constant velocity and constant speed) on perceived distance between the red and blue dots. As with the main data, we also fit a Linear Mixed Effects model to the responses, using an REML criterion with a Nelder-Mead optimizer, participant as a random effect, condition as a categorical fixed effect and frame movement amplitude as a numeric fixed effect with a standard linear link function. These analyses do not add anything substantive to the results as plotted and so are not included in the main text. They are available with the raw data on the OSF site.

## Data and Code Availability

The raw data, summary statistics, ANOVAs and Linear Mixed Effects analyses, as well as the code and the 2 movies are available at OSF (https://osf.io/y5xrn/wiki/home/).

## Supplemental Materials

### Movies

**Movie 1:**
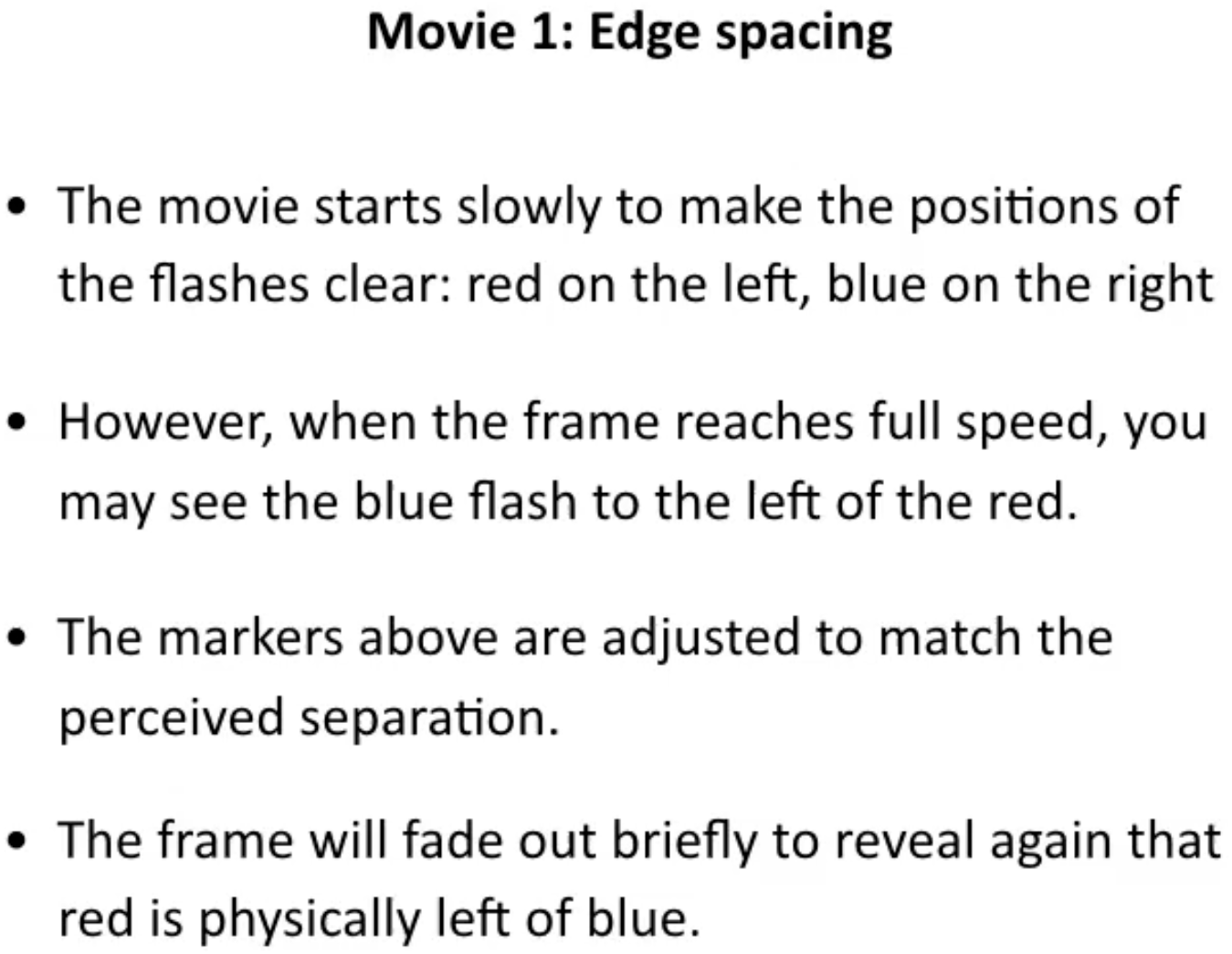
Paradoxical stabilization of a moving frame. The right edge flashes red at the left end of the frame’s travel and the left edge flashes blue at the right end. Participants matched the perceived separation between the flashes with the markers on the upper right. In this example, the frame travel is longer than the frame size so red flash is physically to the left of the blue flash. Nevertheless, blue is seen left of red. Double click on the movie to start or go here https://vimeo.com/492667100 or here https://cavlab.net/Demos/Frame

**Movie 2:**
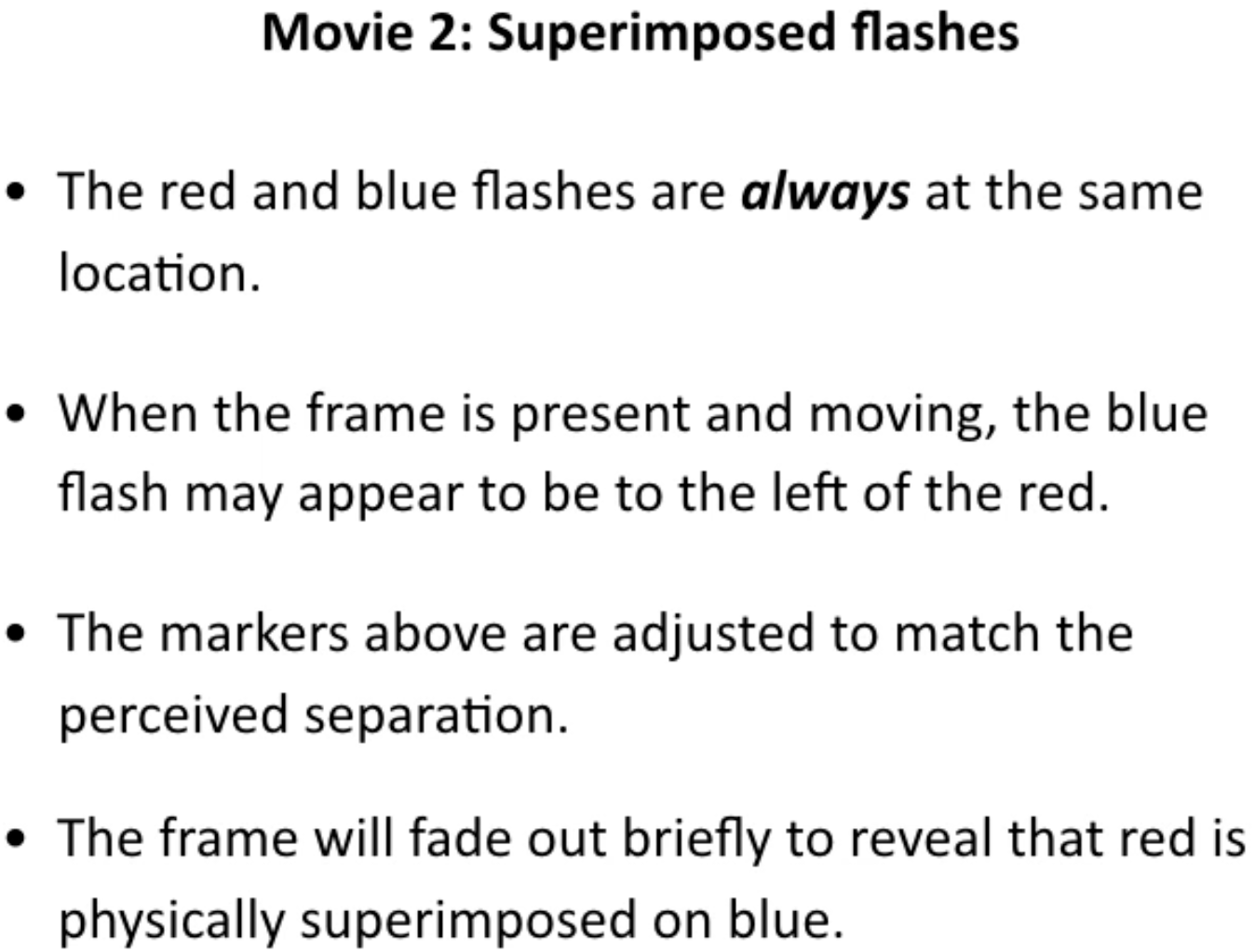
Superimposed flashed probes. The two discs, one red and one blue, always flash at the same location but when the moving frame is visible, they are pulled to the left and right by almost as much as the frame’s displacement. This is not affected much by fixation (try fixating the corners of the movie). Double click on the movie to start or go here https://vimeo.com/492667545 or here https://cavlab.net/Demos/Frame.

## References

1. Johansson, G. (1950). Configurations in the perception of velocity. Acta Psychologica, 7 25–79.

2. Duncker, K. (1929). Über induzierte Bewegung (Ein Beitrag zur Theorie optisch wahrgenommener Bewegung). Psychologische Forschung, 12, 180–259. Abridged and translated (1938) as “Induced Motion” in Source Book of Gestalt Psychology edited and translated by Ellis, W. D. London: Routledge and Kegan Paul. pp 161–172.

3. Asch, S. E., & Witkin, H. A. (1948). Studies in space orientation: I. Perception of the upright with displaced visual fields. Journal of Experimental Psychology, 38 325–337.

4. Morgan, M., Grant, S., Melmoth, D., & Solomon, J. A. (2015). Tilted frames of reference have similar effects on the perception of gravitational vertical and the planning of vertical saccadic eye movements. Experimental Brain Research, 233(7), 2115–25. doi: 10.1007/s00221-015-4282-0.

5. Roelofs, C. O. (1935). Optische Lokalisation. Archiv fur Augenheilkunde, 109 395–415.

6. Matin, L., & Fox, C. R. (1989). Visually perceived eye level and perceived elevation of objects: linearly additive influences from visual field pitch and from gravity. Vision Research, 29(3), 315–24. doi: 10.1016/0042-6989(89)90080-1.

7. Warren, H. C. (1895). Sensations of rotation. Psychological Review, 2(3), 273–276. doi: 10.1037/h0074437

8. Wallach, H. (1959). The perception of motion. Scientific American, 201 56–60.

9. Agaoglu, M. N., Herzog, M. H., & Öğmen, H. (2015). Field-like interactions between motion-based reference frames. Attention, Perception, & Psychophysics, 77 2082–2097. doi: 10.3758/s13414-015-0890-9

10. Cavanagh, P., Wexler, M., & Anstis, S. (2020). Frame-induced position shifts. Journal of Vision, 20(11):607. doi: 10.1167/jov.20.11.607.

11. Reinhardt-Rutland, A. H. (1988). Induced movement in the visual modality: an overview. Psychological Bulletin, 103 57–71.

12. Nakayama, K., & Tyler, W. (1978). Relative motion induced between stationary lines. Vision Research, 18 1663–1668.

13. Wertheimer, A. H. (1987). Retinal and extraretinal information in movement perception: how to invert the Fihlene illusion. Perception, 16 299–308.

14. Hogendoorn, H. (2020). Motion extrapolation in visual processing: lessons from 25 years of flash-lag debate. Journal of Neuroscience, 40(30), 5698–5705. doi: 10.1523/JNEUROSCI.0275-20.2020

15. Cavanagh, P. & Anstis, S. (2013). The flash grab effect. Vision Research, 91 8–20. doi: 10.1016/j.visres.2013.07.007

16. Sinico, M., Parovel, G., Casco, C., & Anstis, S. (2009). Perceived shrinkage of motion paths. Journal of Experimental Psychology: Human Perception and Performance, 35(4), 948–957. doi: 10.1037/a0014257

17. Wojtach, W. T., Sung, K., Truong, S., & Purves, D. (2008). An empirical explanation of the flash-lag effect. Proceedings of the National Academy of Sciences, 105(42), 15338–16343. doi:10.1073/pnas.0808916105

18. Krekelberg, B., & Lappe, M. (2000). A model of the perceived relative positions of moving objects based upon a slow averaging process. Vision Research, 40 201–215.

19. Anstis, S., & Cavanagh, P. (2017). Moving backgrounds massively change the apparent size, shape, and orientation of flashed test squares. I-Perception, 8(6), 1–4. doi: 10.1177/2041669517737561

20. MacKay, D.M. (1973) Visual stability and voluntary eye movement. In R. Jung (ed.), Handbook of sensory physiology, Vol. VII/3. Heidelberg/New York: Springer Verlag, pp. 307–331.

21. O’Regan, K. (1992). Solving the “real” mysteries of visual perception: the world as an outside memory. Canadian Journal of Psychology, 46(3), 461–488.

22. Helmholtz, H. V. (1867). Handbuch der Physiologischen Optik, 1st Edn. Leipzig: Voss. Translated as (1967). Treatise on physiological optics, Volumes 1 and 2 (J. P. C. Southall, Trans.). New York: Dover.

23. von Holst E, Mittelstaedt H (1950) Das Reafferenzprinzip (Wechselwirkung zwischen Zentralnervensystem und Peripherie). Naturwissenschaften, 37 464–476.

24. Wurtz, R. H. (2008). Neuronal mechanisms of visual stability. Vision Research, 48 2070–2089.

25. Wurtz, R. H. (2018). Corollary Discharge Contributions to Perceptual Continuity Across Saccades. Annual Review Vision Science, 4 215–237. doi: 10.1146/annurev-vision-102016-061207

26. Collins, T., Rolfs, M., Deubel, H., & Cavanagh, P. (2009). Post-saccadic location judgments reveal remapping of saccade targets to foveal locations. Journal of Vision, 9(5):29, 1–9.

27. Deubel, H., Koch, C., & Bridgeman, B. (2010). Landmarks facilitate visual space constancy across saccades and during fixation. Vision Research, 50 249–59

28. Cavanagh, P., Hunt, A., Afraz, A., & Rolfs, M. (2010). Visual stability based on remapping of attention pointers. Trends in Cognitive Sciences, 14 147–153.

29. Murakami, I, & Cavanagh, P. (1998). A jitter aftereffect reveals motion-based stabilization of vision. Nature, 395 798–801.

30. Bridgeman, B. (2010). Space constancy: the rise and fall of perceptual compensation. In R. Nijhawan and B. Khurana (eds.) Space and Time in Perception and Action. Cambridge: Cambridge University Press, pp. 94–108.

31. Mack, A., Bachant, J. (1969). Perceived movement of the afterimage during eye movements. Perception & Psychophysics, 6 379–384. doi: 10.3758/BF03212795

32. Mach, E. (1886). Beitrage zur Analyse der Empfindungen. 1st ed. Jena:Verlag von Gustav Fischer. Translated by Fischer J. 1959. The analysis of sensations. New York:Dover.

33. Stevens, J. K., Emerson, R. C., Gerstein, R. L., Kailos, T., Neufeld, G. R., Nichols, C. W., & Rosenquist, A. C. (1976). Paralysis of the awake human: Visual perceptions. Vision Research, 16 93–98.

34. Ditchburn, R. W., & Ginsborg, B. L. (1952). Vision with a stabilized retinal image. Nature, 170 36–37. dio: 10.1038/170036a0.

35. Sperling, G. & Speelman, R. (1965). Visual spatial localization during object motion, apparent object motion, and image motion produced by eye movements. Journal of the Optical Society of America, 55 1576.

36. MacKay, D. M. (1970). Mislocation of test flashes during saccadic image displacements. Nature, 227(5259), 731–3. doi: 10.1038/227731a0

37. O’Regan, J. K. (1984). Retinal versus extraretinal influences in flash localization during saccadic eye movements in the presence of a visible background. Perception & Psychophysics, 36(1), 1–14.

38. Ostendorf, F., Fischer, C., Gaymard, B., & Ploner, C. J. (2006). Perisaccadic mislocalization without saccadic eye movements. Neuroscience, 137(3), 737–45. doi: 10.1016/j.neuroscience.2005.09.032

39. Honda, H. (1995). Visual mislocalization produced by a rapid image displacement on the retina: examination by means of dichotic presentation of a target and its background scene. Vision Research, 35 3021–3028.

40. Morrone, M. C., Ross, J., & Burr, D. (1997). Apparent position of visual targets during real and simulated saccadic eye movements. Journal of Neuroscience, 17 7941–7953.

